# Aerodynamic performance appears to have shaped the evolution of flight-feather microstructures

**DOI:** 10.64898/2026.09.17.752277

**Authors:** F. Alenius, J. Revstedt, L. C. Johansson

**Affiliations:** Department of Biology, Lund University, Lund, Sweden; Department of Energy Sciences, Lund University, Lund, Sweden

## Abstract

Flight feather structures - shaft, barbs, and barbules - have transformed through evolution. Although size, shape and orientation of these structures likely influence the aerodynamic performance of the feathers, their impact remains unclear. Here we investigated the feather structures’ influence on flow and aerodynamic performance, using computational fluid dynamic modelling, by modifying the morphology of a section of a Jackdaw (*Corvus monedula*) flight feather. We show that the flow over the leading vane is a key factor, and that changing the barb-to-shaft angle from its natural state as well as the trailing vane barb angle, reduces performance, which could explain evolutionary changes from early *Paraves* species. All in all, our results suggest that current feather composition may be close to a local maximum for both aerodynamic efficiency and stability/flow predictability, suggesting evolution of feather structures have been driven by aerodynamic performance, which should have implications for designing bio-inspired technologies.

## 3. Introduction

Flight feathers are extremely complex structures operating in a flow regime where the size and shape of the microstructures composing the feather have potential to affect aerodynamic performance ^1^. From the fossil record we know that the morphology of several of these structures have evolved over time ^2–4^, and that the primary flight feathers of the earliest birds (i.e. *Archaeopteryx lithographica*, 150 Mya) differ from those of modern birds ^2,4^. However, our understanding of how these changes in shape have affected aerodynamics is poorly understood. Neither do we know if there exist alternative shapes that could provide a better aerodynamic performance than existing ones but have failed to evolve due to developmental constraints ^3,5,6^ or conflicting selection pressures with e.g. mechanical performance ^3,5^. Understanding the role and optimal shape of these microstructures is thus important for our understanding of evolution of flight in birds.

Feathers evolved before flight ^7–9^, suggesting the original use of feathers was something other than flight (e.g. insulation or display) ^3,8^. As a result, the evolution of flight feathers may be constrained by the way feathers grow, or the trade-off between the multiple functions of feathers. Flight feathers are constructed with a leading vane, a shaft, and a trailing vane ^2,10^. The vanes are built in a hierarchical manner, where barbs branch off the shaft at an angle and barbules branch off barbs ^2,10^ (Fig 1). The barbules of adjacent barbs overlap and join ^10,11^ and the barbs and barbules together form the aerodynamic surface ^10^. The angles between barbs and shaft differ between the leading and trailing vanes in flight feathers ^2,10^ and the angle on the trailing vane changes significantly from a low angle to a higher angle from archaic to modern birds ^2^. However, feathers develop by growing inside a tube ^8,10^ with the barbs and barbules tightly packed, branching at different angles than in the mature feather ^12^. This means the barbs and barbules, despite being dead keratin structures ^3,12^, need to change shape after their growth has finished. Given that evolution reshapes existing structures ^8^, this suggests that some morphological solutions may not be possible to achieve in feathers, such as barbules forming a solid plane, due to how the feather develops. Structural stability and weight minimization ^13^ provide additional selection pressures that may favor shapes that differ from the shapes generating aerodynamic forces efficiently. Alternatively, since feathers are dead structures ^3,12^, control of aerodynamics needs to be mostly passive, and natural selection may favor aerodynamic stability rather than e.g. peak efficiency. Taken together, we may not expect aerodynamic efficiency to be the sole optimization criteria and, in some aspect, feathers may perform aerodynamically suboptimal. To resolve these issues, we need to explore alternative feather morphologies to test the impact on aerodynamic performance.

**Figure 1.**
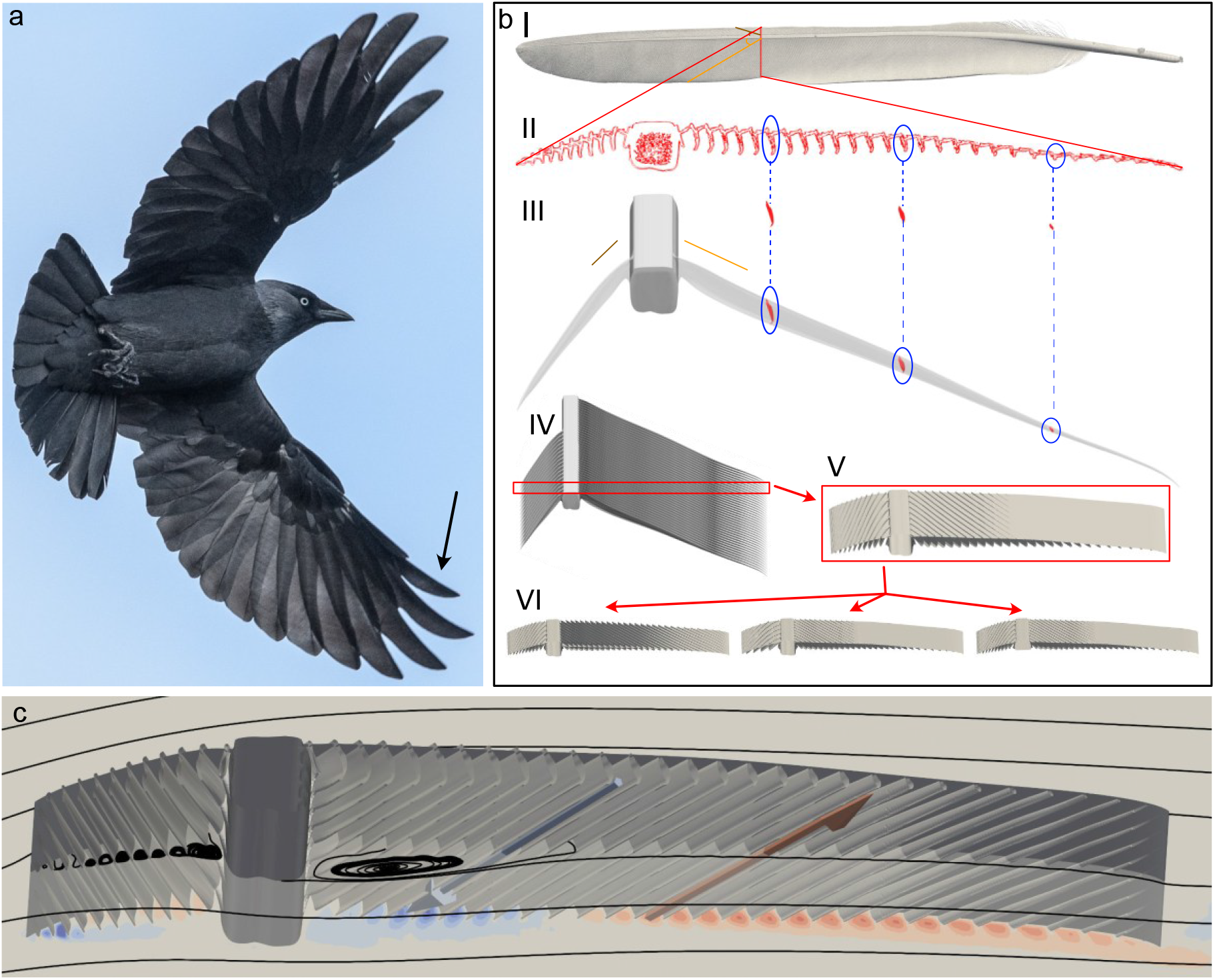
Feather and feather modelling. a) For this study we used the ninth primary feather (marked by an arrow) of a Jackdaw (Corvus monedula). The outer primary feathers spread and form a slotted configuration where our feather is the leading feather. Photo: Arend Vermazeren, used under a Creative Commons license 2.0 b) We CT-scanned a feather (I) and used a cross section (red line) from the part of the feather that function as an independent airfoil, to build the model. (II, III) Cross sections of the barbs were moved into the correct 3D position (as seen in the CT-scan of the full feather) maintaining the angle relative to the shaft (orange and brown lines) and a single barb on each vane was created. (IV) These barbs were then copied and repeated along the shaft. The shaft was created as a tube with the same cross section as the sample cross section. The barbules were modelled as a solid sheet. (V) We then cut the model in the chordwise direction so that the model is self-repeating in the spanwise direction and can create an infinite wing in the spanwise direction. (VI) From the base model we made modifications by moving the barbule plane, adjusting the angle of the barbs or modified the shaft. Image reworked from Alenius et al ^1^ c) The overall flow around the feather model is greatly impacted by the structures of the feather. Between the barbs, the flow rotates close to the opening between the barbs. A larger rotational flow structure occurs behind the shaft in the space between the bottom of the shaft and the bottom of the barbs. Above the rotational structures between the barbs, but underneath the barbule plane, the flow follows the angle of the barbs and depending on the location along the barb, the direction of the flow is either into the paper (blue) or out of the paper (red). As a result, the flow between the barbs is directed forwards at the leading part of both the leading and trailing vane, against the free-stream flow. (Reworked from Alenius et al. ^1^)

Primary flight feathers make up the distal part of a bird’s wing and in many species the outermost primaries separate into a multi-slotted configuration (Figure 1a) as seen in e.g. birds of prey ^14^. When the feathers separate, they each function as individual aerodynamic units (i.e. wings) ^15^, emphasizing aerodynamic performance as a natural selection pressure on the feather microstructures. The feathers in a slotted configuration differ in shape from other flight feathers ^3,10^, making them particularly relevant to study to understand how the microstructures affect aerodynamic performance. The small size of the shaft, barbs and barbules, in combination with wind tunnel and measurement system limitations, makes it difficult/impossible to study the flow near feathers (see Supplement), let alone modify the shape of their microstructures (but see^16^ Tu et al., 2024). In the species used here, the jackdaw (*Corvus monedula*), the chord of the tested feather section is <14 mm and the perpendicular distance between barbs is ~0.23 mm. To be able to investigate the aerodynamic impact of feather microstructures we opted to perform numerical experiments using computational fluid dynamic simulations. This approach allows us to systematically modify the feather structures to compare their individual and relative impact on the aerodynamic performance and deduce if other feather functions may have influenced the evolution of morphology.

## 4. Results and discussion

To test the aerodynamic effect of microstructures, we generated a feather model (FM) based on a CT scan of the ninth primary flight feather of a Jackdaw (Figure 1), with an average vane shape based on the CT scan and four additional surface scanned feathers. The ninth primary is the leading feather of the multi-slotted outer wing (Figure 1a) and the FM represents a section of the feather operating as a wing on its own (Figure 1b). We then modified the model to test the effect on aerodynamic performance of three structural aspects; the location of the barbules along the height of the barbs, the angle of the barbs in relation to the shaft and the height of the shaft (see Material and methods for details).

The first two modifications determine the depth to width ratio of the valleys formed between the barbs, a factor with potential to influence drag and lift at the flow regime of feathers ^17–19^. The third modification investigates one of the main differences in profile shape between feathers and traditional airfoils. We simulated the flow around the models across a range of angles of attack (−2.59°≤α≤10.41°, where α is the angle between the air flow and the feather chord, a line connecting the leading and trailing edge of the model) with the geometry fixed and at gliding flight conditions. Reynolds number (*Re* = *cU*/*v*, where c is the chord length, U the airspeed and ν the kinematic viscosity), indicating the relative importance of inertial to viscous forces, was set to 7000. This represents a flight speed of 8 m/s, which is close to the minimum power speed of Jackdaws ^20^. To evaluate the effect of the morphology, we estimated the lift coefficient (C_l_) (from the force perpendicular to the airflow) and the drag coefficient (C_d_) (from the force aligned with the flow) and the ratio between the lift and drag coefficient (C_l_/C_d_) (a measure of the efficiency of lift generation). We also determined the pitch torque coefficient (C_M_) and the spanwise flow between the barbs (see Material and methods for details).

Compared to the flow around a traditional airfoil, the flow around the FM is more complex due to the intricate feather morphology (Figure 1c). On traditional airfoils, the flow is ordered and follows (remains attached to) the smooth surface at low angles of attack, but can separate (i.e. detach) from the upper surface at moderate to high angles of attack, creating a wake that contributes to more drag ^21–23^. A feather does not have smooth surfaces, with numerous barbs creating ridges on both the dorsal and ventral side, and the shaft protruding into the airflow on both sides (Figure 1b). In the defined spaces between the barbs, on both the dorsal and ventral side, as well as behind the shaft on the ventral side, we found formation of rotational flow structures (i.e. vortices) (Figure 1c). The fact that we find these rotational flow structures suggests that the shape of the morphological structures are important predictors of the performance of the feather model. At the same time, the non-smooth surface of the feather/FM can yield predictability in the flow as separation points become fixed ^1^. On the ventral side we also see spanwise flow between the barbs, which close to the shaft also flows upstream while further away from the shaft flows downstream (Figure 1c). Comparing the performance of the FM to the traditional airfoils and cambered plates, we have previously shown that despite this complex flow the FM produces lift on par with these airfoils for similar *Re* ^1^.

### 4.1. Location of the barbule plane

The depth to width ratio of the valleys between the barbs in the streamwise direction is expected to influence how the flow stays attached to the feather ^24,25^, i.e. affect the separation point and the consequent wake structure above the trailing vane (TV) ^1^. The depth to width ratio is primarily determined by the position of the barbules along the height of the barbs (Figure 2a). Due to the complexity and small dimension of the barbules, we modeled them as a plane (BP), where the position of the plane is given in percentage of the height of the barbs, with 0% being the top of the barb (Figure 2a). We found that altering the position of the BP has a clear impact on the flow (Figure 2e) and performance, influencing both C_l_ and C_d_ of the model. The FM scores high on the integrated lift-to-drag ratio ((*C*_*i*_⁄*C*_*d*_)_*sum*_) and peak C_l_/C_d_((*C*_*i*_⁄*C*_*d*_)_*max*_) relative to most of the modifications (Figure 2b). When moving the BP down on the TV, (*C*_*i*_⁄*C*_*d*_)_*sum*_ and (*C*_*i*_⁄*C*_*d*_)_*max*_ decreased compared to the FM (Figure 2b). However, the highest performing model was with the BP placed the highest 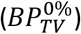. Also note that there seems to be little change between the values of the 25% and 50% position of the BP 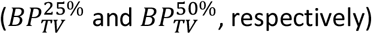 (Figure 2b), suggesting that small changes in position of the BP, in the mid-range, will not have a substantial impact on aerodynamic efficiency or overall airflow.

**Figure 2.**
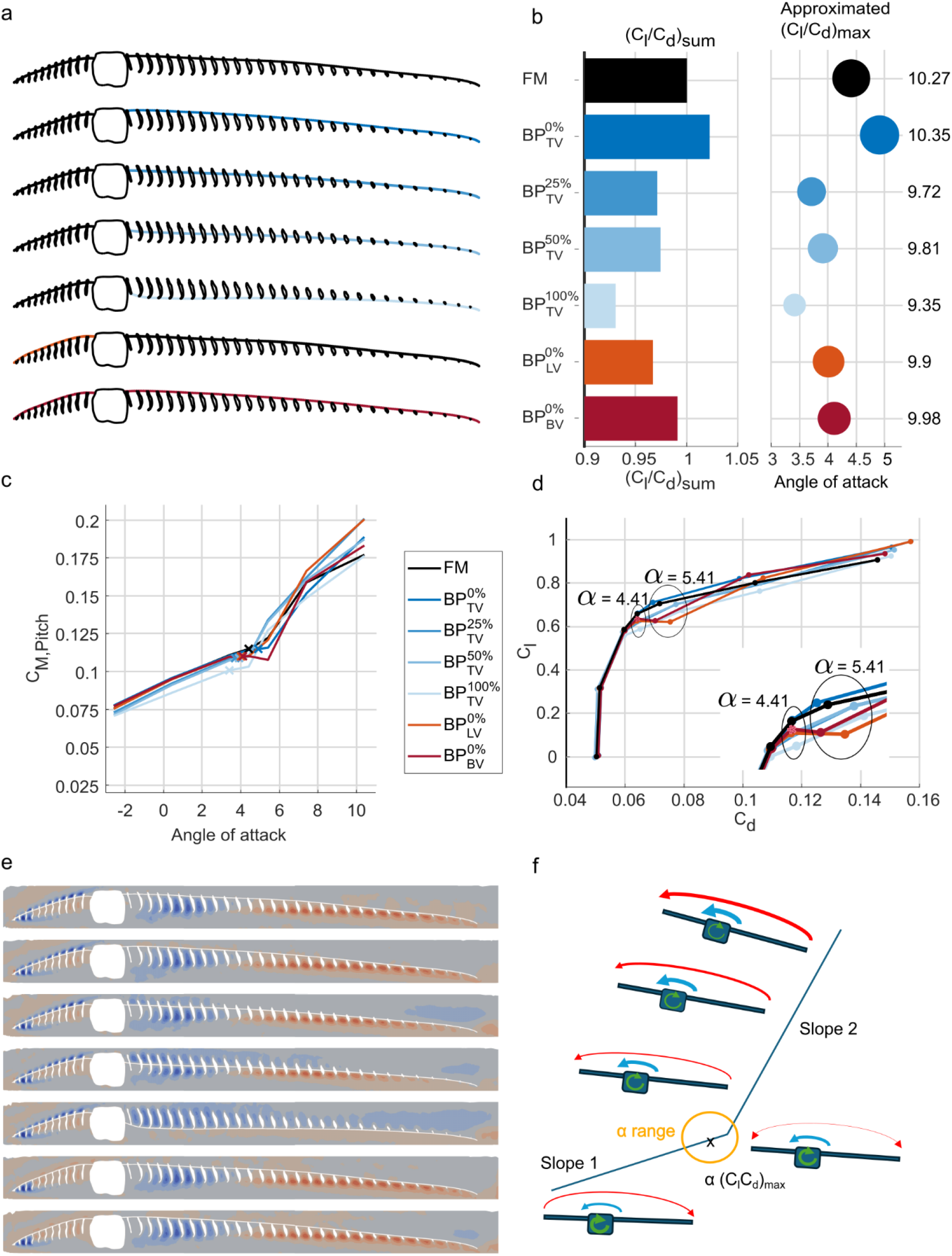
The height of the barbules affects aerodynamic performance. a) Schematic images of the barbule plane modifications on the trailing and leading vane. Colors represent the different modifications given by the abbreviation 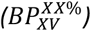, where BP stands for barbule plane, TV stands for trailing vane and LV stands for leading vane, BV stands for both vanes, while the XX% indicates the relative height of the BP along the height of the barbs, where 0% is at the top of the barbs. Feather model (FM) = black, TV modifications = blue, LV modifications = orange and BV modifications = red. b) The integrated lift-to-drag ratio (C_i_⁄C_d_)_sum_ over α relative to FM (bars) and the (C_i_⁄C_d_)_max_ (dot size indicate relative magnitude and number to the right the value) vs α. c) Aerodynamic torque coefficient (C_m_) around the feather shaft center, where positive values give pitch down rotations. The crosses show the α of max C_l_/C_d_ for each of the modifications. d) Polar plots (C_l_ vs C_d_) for the different models with α=4.41° and α=5.41° circled. Note the dip in C_l_ when having a smooth top surface of the LV. Note that 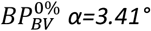, is marked with an asterisk, this is due to a noted disturbance in spanwise flow in front of the foil which is not expected to have a large impact on the overall result. e) Spanwise flow of the different models at α=4.41, direction out of the paper (red) and into the paper (blue). The order is the same as in a and b. f) Schematic image of the proposed mechanism for passive pitch control. Pitch response (red) resulting from the two different (built-in pitch-up green arrow, and aerodynamic, blue arrow) torques and the two different slopes of aerodynamic torque across α. Increasing α increases aerodynamic torque, which tends to reduce α. To achieve low α the feather needs to pitch down, increasing tension in the shaft, which then acts as a spring to pitch up the feather again. The X marks the α of the (C_i_⁄C_d_)_max_ location, the suggested optimum for the feather profile to be at. The yellow circle shows the approximate area where (C_i_⁄C_d_)_max_ occurs for the different modifications.

The lower aerodynamic efficiency associated with lowering the BP on the TV was caused by a combined effect of lower C_l_ and higher C_d_(Figure 2d). The lower C_l_ stems from the altered pressure distribution around the shaft (supplemental Figure S1) and the higher C_d_ was associated with the deeper indentations between the barbs on the dorsal side where the increased unevenness of the surface increases the disturbances in the wake and hence the height of the wake (supplemental Figure S2. We note that the modification has little effect on α≤3.41°, which could be interpreted as the selection pressures for aerodynamic performance mainly have effect on higher angles of attack. However, this study only concerns gliding flight and the responses of the feather could be different for other flight modes (e.g. flapping or maneuvering). Interestingly, the response to a smooth top surface differed between the leading vane (LV) and TV. A smooth top surface on the LV lowered the (*C*_*i*_⁄*C*_*d*_)_*max*_, and (*C*_*i*_⁄*C*_*d*_)_*sum*_ regardless of if the TV was ribbed 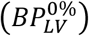 or not 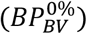 (Figure 2b). For the LV, our results thus suggest that it is favorable to *not* have a smooth top surface. However, this was not mainly caused by a lower C_d_, which has been emphasized in other types of animal foils with non-smooth surfaces (e.g. sharks and dragonfly wings ^26–29^), but instead by a higher C_l_ at α=4.41° and α=5.41° (Figure 2d), similar to the improved performance of tails of swimming sharks caused by the scales ^19^. When investigating C_l_ for all cases where C_l_ is relatively lower than for the FM at α=5.41°, we found that the simulations take longer time to converge, which may be caused by a mode shift in the flow, from a laminar to a transitional or turbulent mode, occurring in this α-range. The shape of the C_l_-curve shows a dip in C_l_, where the exact position is not fully resolved by the current α resolution. Transitioning across the range of α of the dip will cause fluctuation of the forces (C_l)_, which we suggest is something a bird would benefit from avoiding. The fact that we find that a smooth top surface on the TV improved the aerodynamic performance relative to the FM, while real feathers do not have an entirely smooth top surface suggests the location of the BP is influenced by other factors than aerodynamic efficiency. Having the BP below the top ridge of the barbs could be a result of developmental constraints e.g. if connected (genetically or otherwise) to the development of the LV where our model finds a non-smooth surface beneficial. Alternatively, the barbules are delicate structures forming the aerodynamic surface, making them important to protect. Barbule ware would increase leakiness of the surface, likely lowering lift ^30^, and result in a need for more frequent feather molt, which is a costly process ^3^ that may also limit the ability of the bird to fly ^3^. When the wings are folding and extending, during each wingbeat, the feathers slide across each other and allowing barbs to protrude above the barbules could have a protective function reducing barbule ware.

In addition to efficiency and force production, we looked at how the stability of the feather models might be affected by aerodynamic pitch torque. For the FM there are two distinct slopes of how the pitch torque coefficient (C_M_) changes across α, with a lower slope for low α and a higher slope for high α (Fig 2 c, f). The higher the C_M_ the more the feather will pitch nose down and the incline of the C_M_ slopes with α determines the pitch down reaction of the FM. Since too high α may, in addition to lower C_l_/C_d_, result in unsteady flow with fluctuations in forces ^1^ it is important to avoid these α and a steep slope in C_M_ could be beneficial in reducing α. If the feather is assumed to have a rotational resistance in the shaft and an initially high α (as suggested by the inherit pitch-up twist along the shaft, supplemental Figure S3), high torques at high α would drive the feather towards lower α. Low aerodynamic torques at low α, on the other hand, would allow the feather to spring back to a higher α (Figure 2f) in a controlled manner, suggesting that two slopes in C_M_ found in our model may have a passive stabilizing effect on the α used by the feather. Such a passive pitch control could result in a dynamic twist of the feather, something that has been shown to improve efficiency of force production in flapping flight ^31^. With different slopes for different ranges of α it would then be favorable to have the break in C_M_ slopes close to an α that maximize C_l_/C_d_, which is also what we find (Figure 2c). From this reasoning we would predict that the aerodynamic torque and the torsional resistance of the feather should balance at an α close to the (*C*_*i*_⁄*C*_*d*_)_*max*_, although gross feather morphology may complicate this reasoning and require more complex models to resolve. The slopes for all BP modifications are similar to the FM slopes, however, there are differences at which α the slopes change incline (Figure 2c) and where (*C*_*i*_⁄*C*_*d*_)_*max*_ occurs. The lower the BP is on the barb, the lower the α of (*C*_*i*_⁄*C*_*d*_)_*max*_(Figure 2b). The smooth LV modification either lowers the α of the slope shift 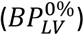 or creates an inconsistency (i.e. a non-linear response) in the lower slope curve 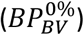. Extending our results to real feathers suggests that since it is unknown where the balance between the aerodynamic torque and resistance of the built-in twist occurs, the changes in torque behavior (both at what α the incline changes and the α of (*C*_*i*_⁄*C*_*d*_)_*max*_) could cause a loss in predictability or non-continuous changes in force production. In the worst case it could introduce sustained pitch oscillation. Such unpredictability could require an active motion of the wing to mitigate the effects, instead of an otherwise passive control of the feather. Given that the behavior is predicted to depend on the mechanical properties of the feather and it is unknown how these C_M_(α)-slopes and location of the shift in slopes are affected by, e.g., wear and tear of a feather that changes over time, but it may be a contributing factor to favor molt. Future studies will need to include mechanical modelling to explore how the fluid structure interactions affect the suggested mechanism and full feather models to determine how this mechanism may interact with other mechanisms, e.g. related to feather sweep^32^.

### 4.2. Angle of the barbs

The barb angle relative to the shaft is a characteristic of primary flight feathers that has been shown to change throughout evolution ^2,8^. The angles differ between the leading and trailing vane and also between feathers in multi-slotted configuration and other feathers ^2^. Here we both increased and decreased the angles on either the LV or TV (Figure 3a) to test if they have any effect on performance. Two of the modifications, increasing the TV angle by ten degrees 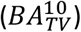 and decreasing the LV angle by five degrees 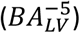 fall inside the morphospace modern birds occupy ^2^, whilst the other two alterations fall outside (Figure 3f).

**Figure 3.**
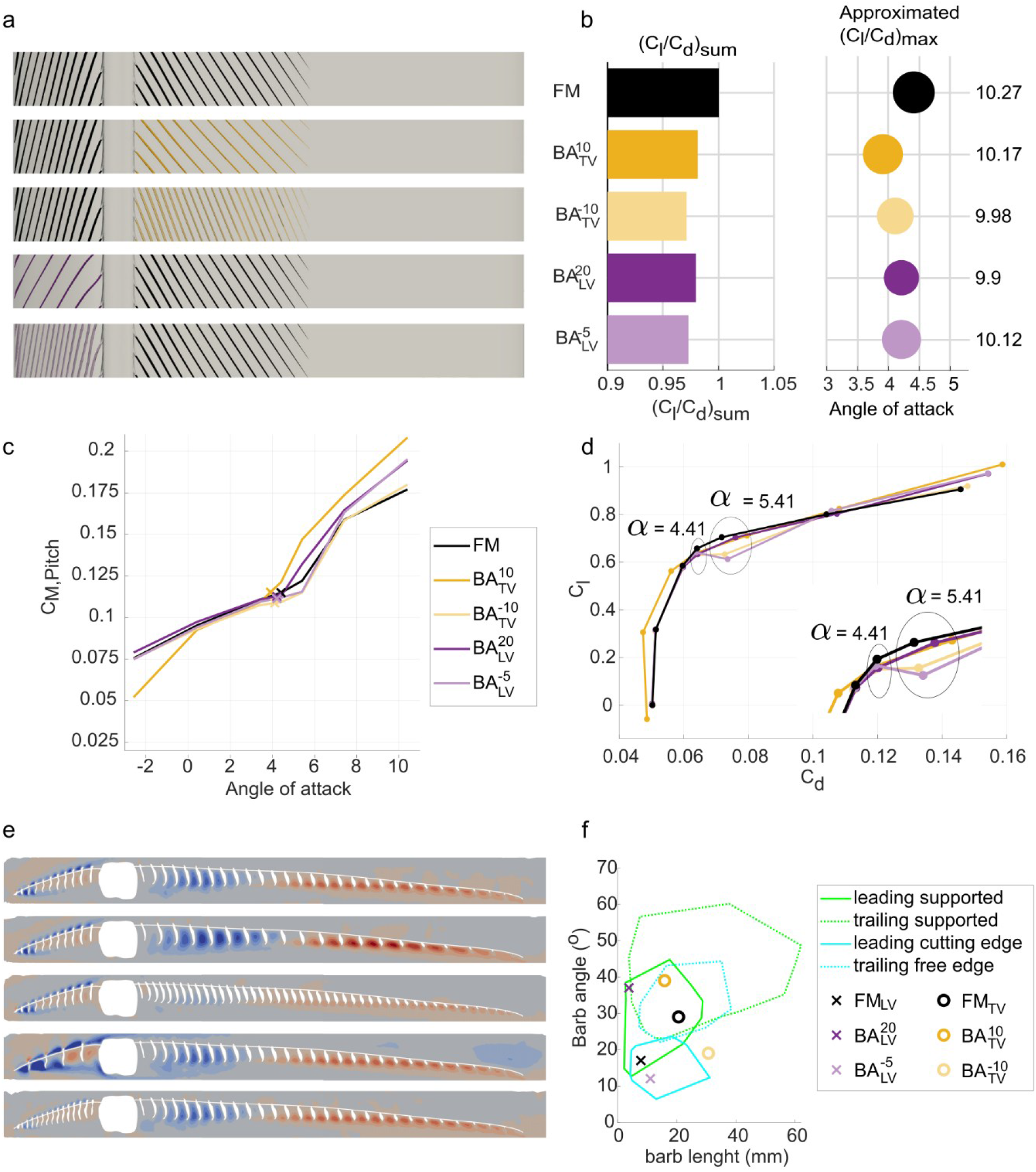
Changing barb angle relative to the shaft lowers aerodynamic performance. a) Schematic images of the barb angle modifications on the trailing and leading vane. Colors represent the different modifications given by the abbreviation 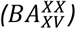, where BA stands for barbule angle, TV stands for trailing vane and LV stands for leading vane and the XX indicates the change in the barb angle (in degrees) relative the original angle. Feather model (FM) = black, TV modifications = yellow, LV modifications = purple. b) The integrated lift-to-drag ratio over α (C_i_⁄C_d_)_sum_ relative to the FM (bars) and the (C_i_⁄C_d_)_max_ (dot size indicate relative magnitude and number to the right the value) vs α. c) Aerodynamic torque coefficient (C_m_) around the feather shaft center, where positive values give pitch down rotations. The crosses show the angle of attack of max C_l_/C_d_ for each of the modifications. d) Polar plot of C_l_ vs C_d_ for the different models with α=4.41° and α=5.41° circled. e) Spanwise flow of the different models at α=4.41, direction out of the paper (red) and into the paper (blue). The order is the same as in a and b. f) Morphospace of the range of barb angle vs barb length of extant birds and feather model (FM) modifications. Green areas represent the part of the feather which is supported by adjacent feathers, Figure 1a, and green represents part of the feather which is not supported (Reworked from Feo et al. ^2^). Circles mark the model and modifications on the trailing edge and crosses mark the model and modifications on the leading edge in the current study.

For all modifications of the barb angles, on both the LV and TV, we found worse aerodynamic efficiency ((*C*_*i*_⁄*C*_*d*_)_*max*_ and (*C*_*i*_⁄*C*_*d*_)_*sum*_) than for the FM (Figure 3b). This can be traced to lower C_l_ and worse C_d_ at mid-range α (Figure 3d). Altering the barb angle while keeping the distance between barbs along the shaft and the vane width constant, changes the perpendicular distance between the barbs. The results show that this influenced the air moving between the barbs, where a larger distance between the barbs result in higher velocity in the spanwise direction, which increases the flow against the free stream direction on the front part of the ventral side of the TV (Figure 3e). The barb angle also affects the chordwise location where the streamwise flow between the barbs, on the ventral side, switches direction (Figure 3e). Decreasing the angle on the TV moved the switch point towards the leading edge while increasing the angle, on both vanes, moved the switch point towards the trailing edge. The barb angle alterations also impacted the C_M_ slopes (Figure 3c), where for α≤3.41° the slopes are very similar except for 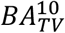, which also have a higher α_zero lift_ than the FM and the other barb angle modifications. At α>3.41° the deviations of the slopes, compared to FM, are larger than for α≤3.41° which is also noted in C_l_ vs C_d_ curves (Figure 3 d). It is notable that the location of the α of (*C*_*i*_⁄*C*_*d*_)_*max*_ remains the same as for the FM, in comparison to the BP modifications that had a larger range.

None of the barb angle modifications give better aerodynamic efficiency than the FM (Figure 3b), suggesting that the barb angles are consistent with optimization by natural selection for aerodynamic performance. In addition to lowering aerodynamic efficiency, several of the tested modifications lead to a significant dip in C_l_ as well as an increase in C_d_, for the angles of attack above peak C_l_/C_d_(α=5.41°). Increasing the BA with 20° on LV 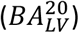, however, did not result in a significant dip in C_l_. This configuration is outside the range of angles for free leading-edge vanes (where free edges do not have support from adjacent feathers ^2^) in modern flight feathers, but fall into the range of supported leading edge vanes (fig 3f). In addition to lowering aerodynamic performance, a large barb angle may lessen the rigidity of the vane ^10^. A rigid leading vane may be needed to uphold an aerodynamic surface, and the larger barb angle also increases the distance between the barbs which puts greater strain on the barbules. The stress on the barbules could be remedied by changing the distance between the barbs, but this could potentially affect the development/growth of the feather and its weight. Increasing the angle on the TV 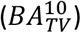 does not generate a dip in C_l_, but a higher C_d_ and thus a lower efficiency, while decreasing the angle 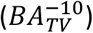 results in a dip in C_l_ at α=5.41° and lower aerodynamic efficiency (Figure 3b). This latter result is interesting given the *BA*_*TV*_ of early birds (e.g. *Archaeopteryx*) have been found to be lower than generally found in modern birds (Figure 3f)^2^. Our results thus suggest that the flight feathers of *Archaeopteryx* may have had lower aerodynamic efficiency than those of modern birds, at least if working in a slotted configuration. Our results consequently give a potential mechanistic explanation for the evolutionary change towards a larger barb angle on the TV previously noted in birds ^2^.

### 4.3. Shaft height

The shaft of the feather protrudes below the lower extension of the barbs of the feather (Figure 4a). We have previously shown that despite this, the FM generates lift on par with airfoils and cambered plates at similar *Re* ^1^. Here we tested the effect of the shaft by decreasing the height of the shaft on the ventral side, to restrict its lower surface to the same level as the lower edge of the barbs, whilst keeping the other relationships (barb angles and barbule plane height) the same as the FM (Figure 4a). This modification altered the airflow below the shaft model significantly, by removing the blockage of the airflow path caused by the shaft, and by extension altering the spanwise flow between the barbs on the TV (Figure 4e). The main effect of the latter is a change in direction of the spanwise, and consequently also the streamwise, flow between the barbs in the first half of the TV (Figure 4e). Furthermore, for α>0°, C_l_ is higher than for the FM except for a dip at α=5.41° (Figure 4d). We also note that α_zero lift_ is increased, which lowers the operational range of the model whilst C_d_ is maintained except for at α<-2° and α>10°, where it is significantly higher than the FM. The change in α_zero lift_ affects the C_M_ for angles below α_zero lift,_ and the C_M_ has three slopes compared to the two slopes for the FM (Figure 4c). Excluding the slope at the lowest α, the other two slopes are similar to those of FM. The dip in C_l_ at α=5.41° occurs above (*C*_*i*_⁄*C*_*d*_)_*max*_ and creates a slight alteration in the C_M_ slope, as well as in the α for (*C*_*i*_⁄*C*_*d*_)_*max*_. This dip in the slope could prevent the passive motion to correct the α to the peak C_l_/C_d_ position described above. Despite this seemingly negative effect, there is a general improvement in aerodynamic efficiency (Figure 4b), since the overall C_l_ is higher than for the FM and C_d_ is not affected. This indicates that these two aerodynamic properties may have to be separated when evaluating overall efficiency. The improved efficiency and largely maintained pitch characteristics, when lowering the height of the shaft, suggests that shaft height has not primarily evolved for aerodynamic function. Instead, we propose structural properties, which should be positively impacted by increased cross section area or second moment of area of the shaft (Figure 4f) ^13,33^, to have been a likely driving force.

**Figure 4.**
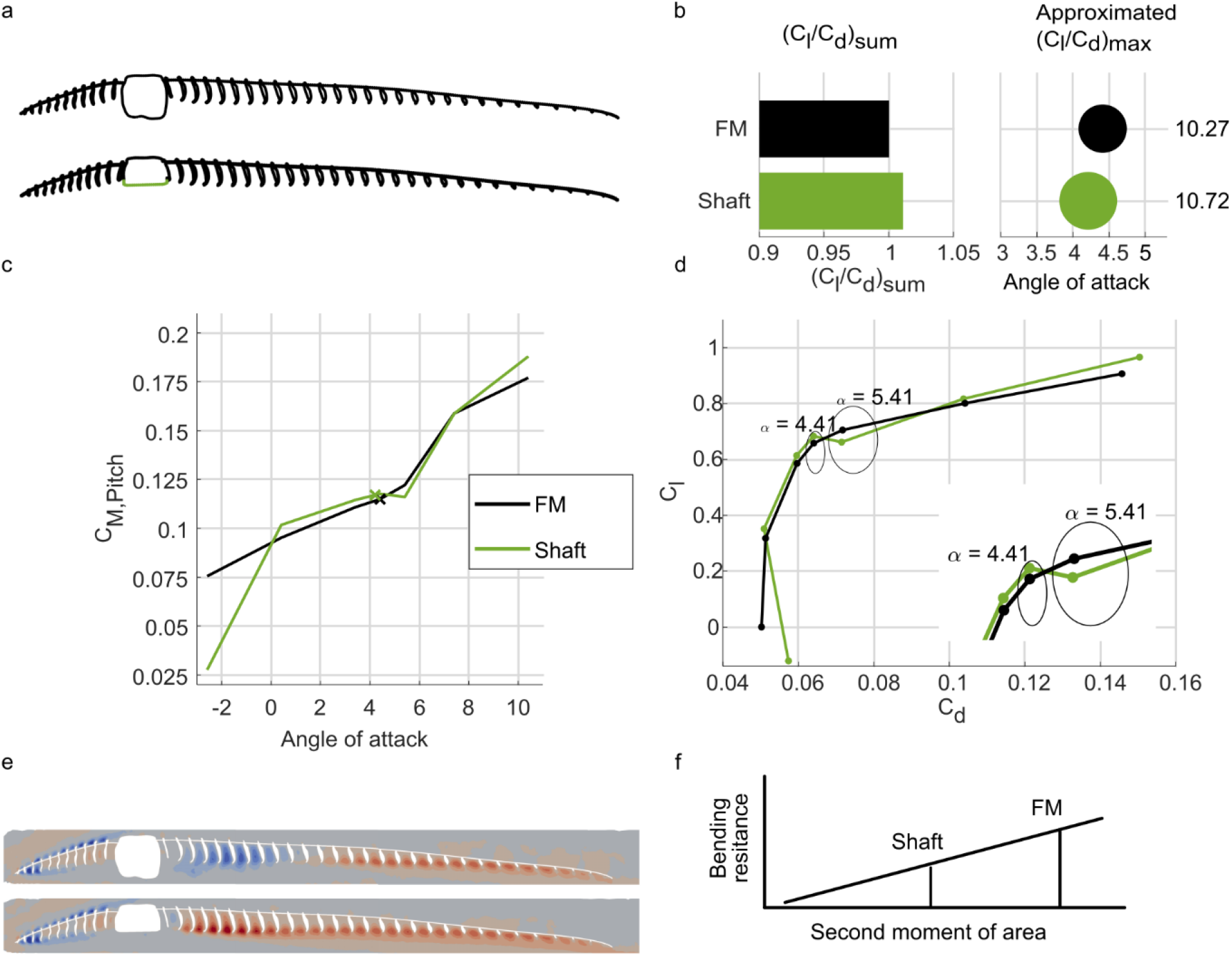
The shaft decreases the aerodynamic efficiency of the feather. a) Schematic images of the shaft modification. Colors represent the different modifications, feather model (FM) = black, Shaft = green. b) The integrated lift-to-drag ratio over α (C_i_⁄C_d_)_sum_ relative to FM (bars) and the (C_i_⁄C_d_)_max_(dot size indicate relative magnitude and number to the right the value) vs α. c) Aerodynamic torque coefficient around the feather shaft center, where positive values give pitch down rotations. The crosses show the angle of attack of max C_l_/C_d_ for each of the modifications. d) Polar plot of C_l_ and C_d_ for the different models with α=4.41° and α=5.41° circled. e) Spanwise flow of the different models at α=4.41, direction out of the paper (red) and into the paper (blue). The order is the same as in a and b. f) Schematic graph showing how bending resistance is expected to vary with the second moment of area of the shaft cross section.

## 5. Conclusions

In this study we have modified several aspects of feather microstructure shape, known to either vary little between species ^2,34^ or to have changed through the evolution of flight feathers ^2,34^. We argue that our findings suggest that the original feather model is close to a local maximum from an aerodynamic efficiency and stability or force predictability perspective. Overall, model modifications on the leading vane show an aerodynamic disadvantage as well as worse stability features when going beyond the angle of attack of peak performance (Figure 2 and Figure 3) compared to the original feather model. This reduction in performance could be a reason why early representatives of *Paraves* already had evolved barb angles similar to extant birds ^2,34^, and that these angles since then have remained. The higher performance of the model with “ribbed”, compared to smooth, leading vane top surface, suggests that the microflow as well as overall flow over the leading edge may have been important factors shaping feathers through the evolution of birds. Model changes of the height of the barbule plane on the trailing vane do not have as drastic consequences on performance as on the leading vane and show more continuous degradation of performance. However, when changing the barb angle on the trailing vane we see reduced aerodynamic efficiency ((C_l_/C_d_)_max_) compared to the feather model. Given that extinct birds, including *Archaeopteryx*, had a much lower barb angle than extant birds, we propose that this evolutionary change in shape may have been driven by aerodynamic performance. We also note that most of our modifications only differ in performance from the FM at α>3.41°. This suggests that any aerodynamic selection pressure mainly applies to high angles of attack, at least for gliding flight. The dip in C_l_ at *α* directly above that of (C_l_/C_d_)_max_ that we found for many of the shape alterations, which also affected the pitch torque curves, indicates a potential predictability issue for a bird. This behavior means that an increased angle of attack does not result in increased or at least maintained lift, but in decreased lift. Consequently, since flying conditions can change rapidly (e.g. from wind gusts), a smooth passive adaptation to these new conditions would be favorable, but the dip in C_l_ suggests this will not be the case. Our results thus suggest that small changes in feather morphology may reduce flight stability or result in reduced aerodynamic efficiency. In cases where our results instead show an increased efficiency when modifying the feather model (e.g. Figure 4), we find potential links to negative impact on other feather functions, such as structural stability or durability.

This study represents a first attempt to evaluate the complex microstructures of feathers and as such relies on simplified models and relative comparisons. Overall, the FM was found to have 1-7 percentage points better performance than most modified models, and only the 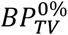 and Shaft modifications improved aerodynamic performance compared to the FM. This aligns with a conclusion that the feather microstructures, at least of our Jackdaws, have been under natural selection for aerodynamic performance. Although the efficiency gain of each morphological change may seem limited, the high energetic demand and the high ecological value of flight for birds is expected to increase the fitness value and allow natural selection to have the structures evolve in large populations^35^. One of the most important future improvements of the model would be to extend it to a full feather and incorporate feather flexibility, or fluid-structural interaction, to allow for passive deformations of the feather. This would allow both testing the effect of mechanical properties and the effect of sweep and twist of the feather, factors that may interact with the effects of the microstructures studied here. Looking beyond the field of bird flight, the aerodynamic function of microstructures highlighted by our findings should be of interest when constructing low Re wings or propellers. Particularly for flexible blades with a torque profile providing self-stabilizing function operating in similar conditions as feathers.

## 6. Material and Methods

We created a feather model (FM) based on a 10 µm/voxel resolution CT-scan of the ninth primary feather of a jackdaw (*Corvus monedula*) (Figure 1a) using the ZEISS Xradia XRM520 Submicron Imaging System (http://www.zeiss.de/) at the 4D Imaging Laboratory, Division of Solid Mechanics, Lund University. This feather was selected since it forms the leading edge, in a slotted configuration, of the outer part of the wing and the distal half of the feather acts as an independent airfoil during flight. Using this scan together with surface scans of four additional feathers (see ^1^ for details), we created a high-resolution CAD-model of a feather section from a cross section where the feather is not supported by its neighbors (Figure 1b). The cross section of the shaft was extruded to create the span of the model. Then, one barb on the leading vane and one barb on the trailing vane were generated by moving the barb cross sections to their 3D position of a single barb on each vane in the full feather scan. These barbs were then copied along the span of the FM, enough times to cover the selected span width. Due to their complexity, we modelled the barbules as a solid plane (BP) instead of the tiny, independent, interlocking structures they are. The plane was placed at the location where the distal barbule attaches to the barb, the barbules do not have the same attachment point along the barb over its length and therefore, the distance from the BP to the top of the barbs is not the same along the chord (c). At x/c > 0.5 the barbule plane and the dorsal point of the barbs are co-located and the top surface is therefore smooth downstream of this point. The width (i.e. the span) of the model was selected so that it could be seamlessly repeated, and the final model had a chord of 13.2 mm and a width of 4 mm. For the leading vane barbs, the angle relative to the shaft was 17° and the spacing 0.8 mm along the shaft. The barbs on the trailing vane have an angle of 29° and a spacing of 0.5 mm along the shaft. The CAD-models were created using SolidWorks2020 (Dassault systems Waltham, Ma, United States).

The original FM model was created by scanning feathers from deceased birds, obtained through the city of Malmö, Sweden, from naturally deceased animals and pest control. No ethical permit was required, since no animals were euthanized for the purpose of this study.

### 6.4. Model and modifications

In addition to the original FM, we created modifications of the model to be able to test the aerodynamic effects of the microstructures. The modifications, either to the shaft, barbs or barbule plane, were created with the same methodology as the FM. The modifications can be divided into three general groups; position of the barbule plane (BP), angle of the barbs (BA) and height of the shaft (Figure 2-4).

#### 6.4.1. Barbule Plane position

The BP position, (Figure 2a), was set as the relative distance from the top of the barbs, with values presented in percentage of barb height. A total of 6 configurations were modeled with a moved BP.

Four modifications with the BP position at 0, 25, 50 and 100% on the trailing vane 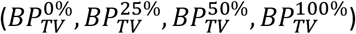, one case with the BP position of 0% on the leading vane 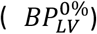 and one case where the BP position of 0% on both the leading and trailing vane simultaneously 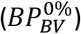.

#### 6.4.2. Barb angle

The BA is the angle (in degrees) between the shaft and the barbule, (Figure 3a). Changing the angle of the barb and keeping the same barb length, would change the streamwise length of the vane and thereby the chord of the model as well as the relative position of the shaft along the chord. To maintain the same model chord and *Re* as the FM, the change in BA was achieved by moving the cross sections of the barbs from the original CT scan in the spanwise direction to achieve the new angle. We modelled a total of 4 configurations with altered BA. Two cases where the BA was modified by −5 and +20 degrees relative to the FM on the LV 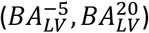, and two cases where the BA was modified by −10 and +10 degrees relative to the FM on the TV 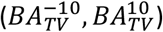.

#### 6.4.3. Shaft height

The shaft protrudes into the flow on the bottom side of the feather (Figure 1b and Figure 4a) and we removed the bottom part of the shaft to create an uninterrupted line between the leading and trailing edge of the model. Only one case with a modification of the shaft has been created (Shaft).

### 6.5. Mesh and simulations

#### 6.5.4. Mesh and domain

We used Hypermesh 2021 (Altair, Troy MI, United States) to create the surface and volume meshes. The outer simulation domain extended 5c above and below, 5c in front, and 10c behind the center of the shaft. The inlet, outlet, top and bottom mesh surface size was set to 6 % of the chord length. We used a refinement box with walls, 0.5c above and below, 0.75c in front, 1.25c behind the center of the shaft with the mesh surface setting of 3 % of the FM chord on the top, bottom, front and back surface. The surface mesh on the FM was set to have an edge deviation of 0.15-0.38 % of the chord, and the side of the domain was set to have a growth rate of 1.05 in the refinement box and an even growth rate outside the refinement box. The mesh settings is summarized in the supplement and discussed in full in a previous study for further details see ^1^.

#### 6.5.5. Simulation settings

We used OpenFOAM8 (openfoam.org) to run the computational simulations. The boundary conditions on the inlet, outlet, top, and bottom conditions are set to free-stream velocity with the streamwise and vertical velocity to match the angle of attack. The walls of the FM and modifications were set to no-slip wall conditions and the side of the domain was set to cyclicAMI, which create an infinitely wide domain allowing the air to move along the barbs uninterrupted. To solve the Navier stokes equations a PISO implicit unsteady simulation was used. Given that the typical gliding speed for jackdaws are approximately 8 m/s ^36^ the chord based Reynolds number will be 7000. At this low Reynolds number the boundary layers are laminar but the wake may be turbulent, it is therefore not suitable to use RANS modelling and any turbulence was instead handled implicitly through the discretization. The diffusion scheme was set to Gauss linear corrected (2nd order), and the temporal discretization scheme was set to Euler (1st order) for the first 0.5 seconds due to stability issues and then switched to backwards (2nd order) whilst the convection scheme is a second order central scheme blended with a certain amount of upwinding (LimitedLinearV 0.1) for stability reasons. With these settings on time and space, turbulence should be captured when present. The model was spatially scaled by a factor 10 and the velocity was therefore scaled down to 0.8 m/s to keep the chord based *Re* of 7000. The simulation validation is summarized in the supplement and discussed in full previously, for further details see^1^.

### 6.6. Extraction of results

All results presented are averages over a time period of 1.5 seconds of the simulations, which equals Ut/c = 9.1. Presented 2D data was extracted at a span position of 40 % of the model if nothing else is stated. Data was further processed in Paraview (paraview.org) or Matlab r2022b (Mathworks, Natick MA, USA). Lift (L) and drag (D) coefficients, C_l_ and C_d_, were calculated according to Eq. 1 and Eq. 2, respectively, using data from OpenFOAM8 monitors, where the lift direction is set perpendicular to, and the drag direction is set parallel to the free-stream flow.

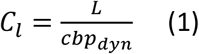

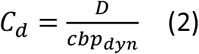

where, *c* is the chord, *b* the span of the model and *p*_dyn_ the dynamic pressure (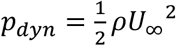, where ρ is the air density and *U*_∞_ is the free stream velocity).

Since the feather is connected to the rest of the bird wing through the shaft, the center of the shaft was used as a center point to determine the torque (M) using the distribution of the aerodynamic forces over the entire surface of the model (Eq. 3).

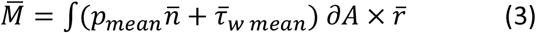

where *p*_mean_ is the time average pressure, 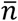 is the surface normal vector, 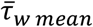 is the time average wall shear stress and r is the distance to the center of the shaft.

The pitch torque (M_Pitch_) is defined as the spanwise torque component of 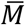 around the shaft center rotating the feather in a counter clockwise motion, Figure 5. Hence the pitch torque coefficient is defined according to Eq. 4.

**Figure 5.**
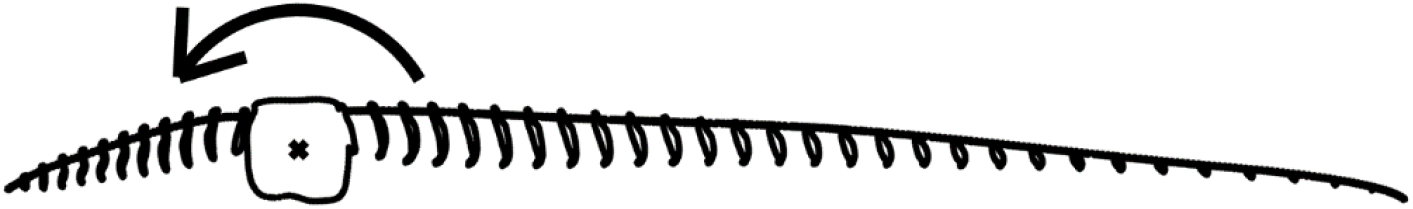
Definition of the positive pitch moment direction.

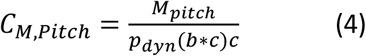

where *c* is the chord, *b* the span of the model and *p*_dyn_ the dynamic pressure.

As a measure of aerodynamic efficiency, we used the lift-to-drag ratio, C_l_/C_d_. To estimate the max value of C_l_/C_d_((*C*_*i*_⁄*C*_*d*_)_*max*_) we used a modified Akima piecewise cubic Hermite interpolation (Matlab function makima) with a resolution of α of 0.1. From the interpolated curve we estimated the max value of C_l_/C_d_ and the associated α. In the figures the (*C*_*i*_⁄*C*_*d*_)_*max*_ was represented as a circle (e.g. Fig 2b) with the radius estimated according to Eq 5.

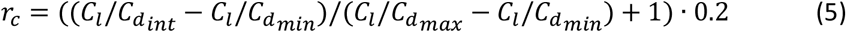

where r_c_ is the radius, (*C*_*i*_⁄*C*_*d*_)_*int*_ the Cl/Cd of the current model, (*C*_*i*_⁄*C*_*d*_)_*min*_ the minimum (*C*_*i*_⁄*C*_*d*_)_*max*_ of all the models and (*C*_*i*_⁄*C*_*d*_)_*max*_ the maximum (*C*_*i*_⁄*C*_*d*_)_*max*_ of all models. The radius was then normalized to have a value between 0.2-0.4.

We evaluated the integrated lift-to-drag ratio, (*C*_*i*_⁄*C*_*d*_)_*sum*_, a measure of the overall performance of the model across a range of angles of attack, by calculating the area under the C_l_/C_d_ curve for positive values of C_l_/C_d_ according to Eq. 6. For comparisons the values were normalized by the value of FM (91.8).

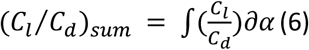

We determined the length of the barbs according to Eq. 7.

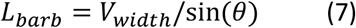

where L_barb_ is the length of the barb in mm, V_width_ is the width of the vane (either leading or trailing) and θ is the angle between the shaft and the barb.

Data on ranges of barb angle versus barb length from extant birds (Figure 3f) has been extracted using plotdigitizer (plotdigitizer.com) from Feo et al. ^2^.

## 7. Acknowledgements

Funding has been provided by the Swedish Research Council grant no 2017-03890 and 2022-02850 to LCJ. The project also received support from eSSENCE grant 3.3 to LCJ, JR and Kent Persson. The computations were enabled by resources at LUNARC Aurora and NSC Tetralith provided by the Swedish National Infrastructure for Computing (SNIC) partially funded by the Swedish Research Council through grant agreement no. 2018-05973 and National Academic Infrastructure for Supercomputing in Sweden (NAISS), partially funded by the Swedish Research Council through grant agreement no. 2022-06725. We are grateful to Kent Persson for discussions regarding the planning of the study, to Stephen Hall for providing the CT-scanning and to Arne Hegemann for providing jackdaws. Several students have provided initial tests that helped decide what the final model should look like.

