## Supplement information for "Aerodynamic performance appears to have shaped the evolution of flight-feather microstructures"

### Summary of content

**Material and methods**

**Mesh Sensitivity**

**Validation**

**Figure list**

Figure S1. Pressure coefficients for FM and BP modifications
Figure S2. Dorsal wake for FM and BP modifications
Figure S3. Scanned CT feather

**References**

**Material and methods**

Prior to the current study we conducted a comparison of the performance of the original feather model (FM) to manmade foils (Alenius et al., 2026). As part of that study, we conducted a mesh sensitivity analysis and a numerical validation of the settings used in the modelling. Here we present a brief summary of those results and refer to the original paper for details.

**Mesh Sensitivity**

Mesh sensitivity was determined, using three surface mesh resolutions, one with half the surface resolution we finally used, one with the final mesh resolution and one with twice the surface resolution of the final mesh. The results of the coarse mesh showed the most difference from the other two and the $C_l$ deviation between the final mesh to the high resolution mesh was less than 1 percent for α=0.41° and α=5.41° and 3% for α=4.41°, while the $C_d$ deviation was less 2% for α=0.41° and α=4.41 and less 3% for α=5.41°. Increasing resolution is expected to reduce the variation in the results. In our case the difference between the final mesh and the high-resolution mesh are within a few percent, even at the angles of attack where flow characteristics are most sensitive to resolution.

**Validation**

We considered validation through experimental measurements of the FM to confirm the results. As noted in the Material and methods, the feather structures are tiny relative to the size of the feather, which makes experiments challenging and we did not manage to perform the experiments for several reasons. First, 3D-printing a 10x enlarged version of the model resulted in model deformations during the cooling process (despite adding support structures), so that it no longer corresponded to the CFD-model. In addition, due to the very thin structure towards the trailing edge, the printed model showed flexibility, which the CFD model did not include. Second, the high complexity (distance between barbs was less than 2.5 mm and varying barb angles on the leading and trailing vane) prevented us from performing PIV measurements close to the surface since the cameras could not view inside the valleys between the barbs.

Instead, we performed a validation study of the simulation settings using an Eppler E387 at Re=10000 across angles of attack in the range from -6 to 18 degrees. In addition to the base case, we studied the influence of mesh resolution, discretization scheme and domain size. The mesh resolution was studied with one case with $dx/c=7.81\times10^{-4}$ (half base case size) closest to the surface and one case with a prismatic refinement on the original mesh. Additionally, another convective scheme has been tested, as well as two cases where the domain size was considered, one with doubling the size in length and height directions and one with a doubling in the span-wise direction. The numerical validation study of mesh resolution, numerical discretization, and domain size show a robust response to the changes made, with little deviation to the base case. For validation, experimental data of E387 (McArthur, 2007) was selected, however, afterwords, we noted that this particular data stands out in performance relative to other airfoils (i.e. Cl & Cd deviates by a factor ~1.2, which is close to air density suggesting the data had not been normalized correctly). Simulations on additional profiles, E61 and NACA4403, and computational results of the same profile (Sauvageat, 2016) reinforces this notion. The numerical validation shows our model to be stable and alternative simulations of the same profile (Levy and Seifert, 2009; Winslow et al., 2018) support our conclusion.

### Figures

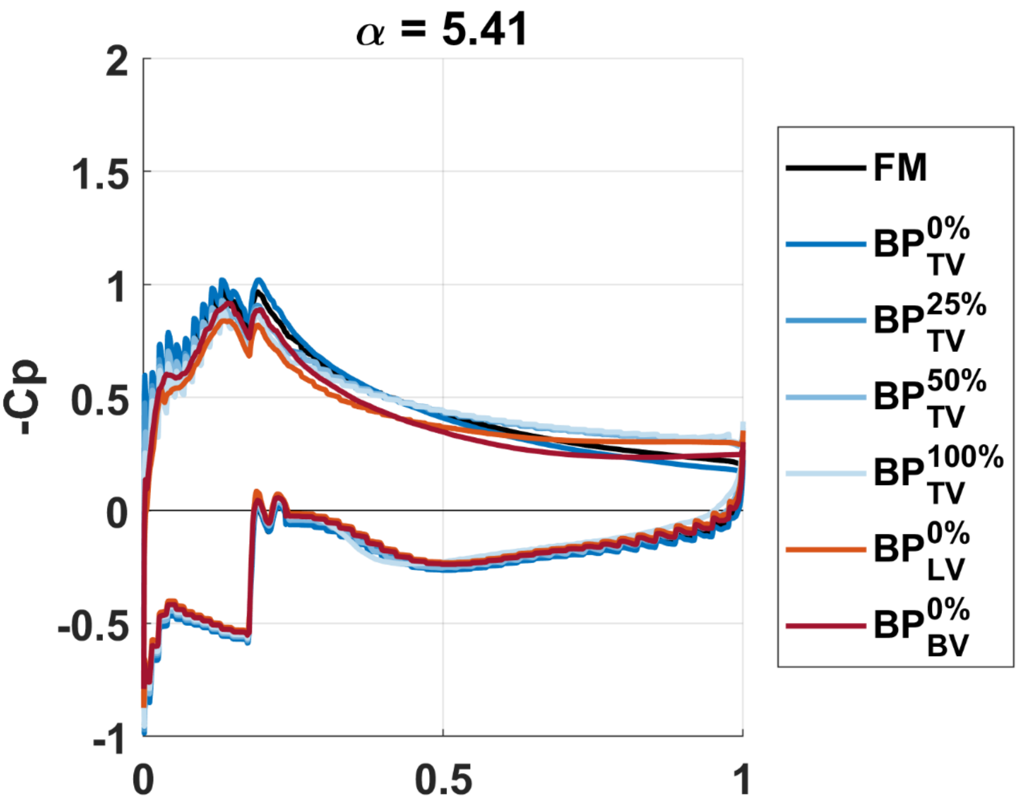


**Figure S1.** Pressure coefficient (Cp) values for the feather model (FM) as well as barbule plane (BP) modifications for α=5.41 at a cross-sectional plane located in the middle of the model. The altered distribution of the pressure on the suction side of the feather (top lines), around the shaft, has an impact which can explain the Cl drop seen in Fig. 2d.

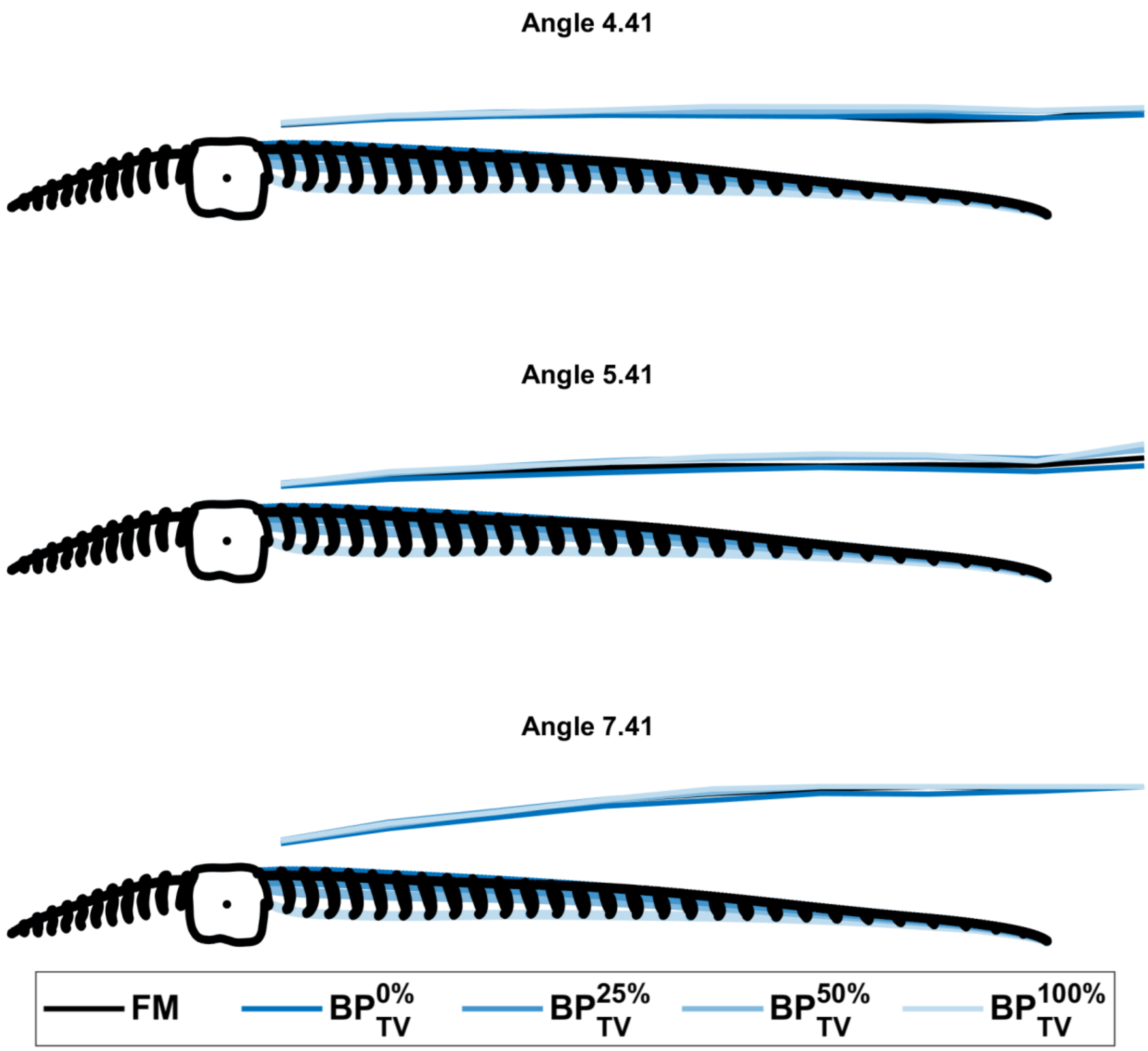


**Figure S2.** Extension of the wake on the dorsal side, above the model. The extension of the wake is defined at the location where the streamwise velocity has reached 95% of the free stream velocity. The further down the barbule plane is, the further the wake extends from the model. The colored lines on the trailing vane represents the different locations of the barbule plane and the responding wake extension line has the corresponding color.

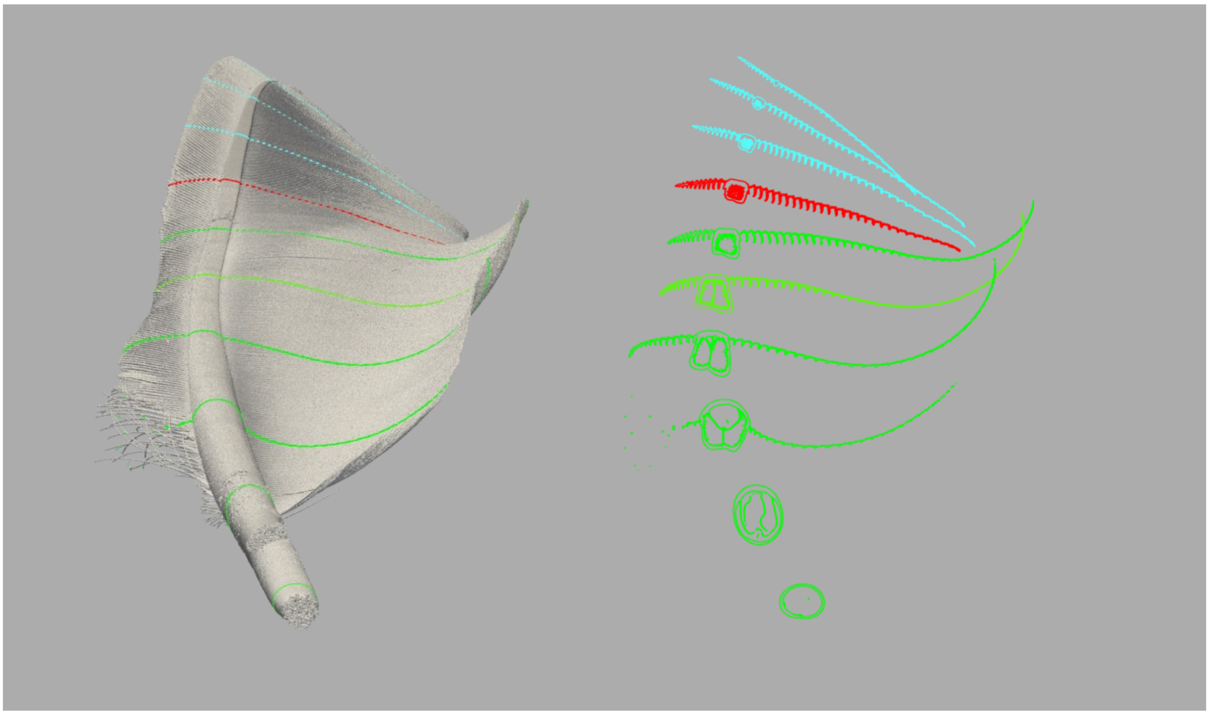

**Figure S3.** CT scanned feather with cross sections marked, green = part of the feather supported by adjacent feathers, red = position of the model and blue = distal part of the feather. Red and blue are in the unsupported part of the feather. As the supported part of the feather ends, a pitch up twist is noticeable. This image is reworked from (Alenius et al., 2025).
